# Natural limits on male mating sustain costly female choice, exaggerated displays, and genetic variation

**DOI:** 10.64898/2026.10.14.759265

**Authors:** Brian Hollis

## Abstract

The persistence of costly female preferences, exaggerated male sexual displays, and the genetic variation underlying those displays remains a central puzzle in evolutionary biology. Classical Fisherian models explain preference-driven display exaggeration but not how costly preference and display can persist without runaway or collapse. These models assume that preferred males remain available regardless of previous mating success, precluding feedback through declining availability. Here I use a gene-centered analysis to show that allelic realized relatedness governs the indirect benefits of choice and that unlimited availability allows relatedness to remain high until genetic variation is lost. Individual-based simulations relaxing this idealization exposed negative feedback: as preference spread and concentrated mating success, preferred males reached capacity and became unavailable. Choosy females became less able to steer indirect fitness benefits toward additional copies of the preference allele, reducing allelic realized relatedness and constraining further evolution of preference and display. This transformed Fisherian runaway into stable exaggeration, allowing costly preference and display to persist for tens of thousands of generations across broad ranges of preference strength, mating capacity, and recombination rate. With autosomal display and a 1% viability cost of preference, stable exaggeration reached 16.5 additive-genetic standard deviations above matched no-preference baselines. Negative feedback generated disruptive selection on display, amplifying additive genetic variation beyond mutation–selection balance by as much as 32-fold, providing a resolution to the lek paradox—the persistence of display variation despite strong sexual selection—without invoking biased mutation or condition dependence. Z linkage of display loci, relevant to birds and other taxa with ZW sex determination, expanded the conditions supporting persistent preference and display, increased exaggeration, and further amplified genetic variation. Once inherent biological limits on male mating success are recognized, Fisherian sexual selection is revealed as a self-regulating form of gene-level cooperation.

## Introduction

Sexual selection has produced some of the most conspicuous traits in nature, including elaborate ornaments and complex courtship displays. Many ornaments and displays arise through female mating preferences, but choosing among potential mates requires time, energy, forgone mating opportunities, or other direct costs (Jennions & Petrie, 1997). Why, then, should choosiness spread and persist? Resolving this question is central to understanding how costly female preferences, exaggerated male displays, and the genetic variation underlying those displays all endure over evolutionary time (Borgia, 1979; Kirkpatrick & Ryan, 1991).

Several mechanisms have been proposed to explain how mating preferences evolve despite costs to females. In some systems, females obtain direct benefits from mate choice, such as resources, parental care, or protection (Jones & Ratterman, 2009). Alternatively, females might obtain indirect genetic benefits by mating with males carrying alleles associated with higher offspring fitness, although empirical support for this “good genes” idea is mixed (Møller & Alatalo, 1999; Prokop et al., 2012). These mechanisms may explain some female preferences, but they do not readily account for preferences for arbitrary male traits. The Fisher process (Fisher, 1930) provides the most influential explanation for such preferences—when females preferring a particular display mate with males expressing that display, their offspring inherit both preference and display, creating a genetic association between the two. This association allows preference alleles to spread through the mating success of display-bearing sons, permitting arbitrary male displays to evolve (Kirkpatrick, 1982; Lande, 1981). Because the Fisher process can in principle explain preferences for any heritable display, some authors have argued that it should be treated as the null model for sexual trait evolution (Prum, 2010), whereas others have suggested that accepting the process without strong empirical support is “methodologically wicked” (Grafen, 1990).

These contrasting views stem partly from a longstanding theoretical problem. Classical models allow runaway evolution, but when female preference is costless they also yield a continuum of neutral preference–display equilibria, with no restoring force maintaining any particular combination of preference and display (Kirkpatrick, 1982; Lande, 1981). Once preference carries a direct cost, preference and display generally decline (Bulmer, 1989; Pomiankowski, 1987), although sufficiently strong genetic associations can instead drive runaway evolution (Hall et al., 2000). Neither outcome explains the stable persistence of costly preference and exaggerated display. Several mechanisms have been invoked to stabilize costly preference and exaggerated display, most notably directionally biased mutation that continually introduces alleles opposing exaggeration (Pomiankowski et al., 1991), although whether biased mutation alone is sufficient depends on how mutations jointly affect ornament size and cost (Dashtbali & Henshaw, 2024). Frequency-dependent mate-sampling costs (Kokko et al., 2015) and evolving mate-search effort (Waffender & Henshaw, 2023) provide two other routes to stability, but the cost of choice vanishes at equilibrium in the former, whereas preference itself carries no direct cost in the latter.

A related but distinct problem is that sexually selected traits often retain unexpectedly high genetic variation despite strong directional selection (Pomiankowski & Møller, 1995). Proposed explanations include host–parasite coevolutionary cycles that maintain variation in resistance signaled by sexual displays (Hamilton & Zuk, 1982) and condition-dependent expression that allows sexual traits to capture genetic variation in overall condition (Rowe & Houle, 1996). Recent theory also shows that mate choice can itself elevate trait variance under some conditions (Xu & Servedio, 2026).

Implicit in most theoretical treatments of Fisherian sexual selection is the assumption that preferred males remain available to females regardless of previous mating success. This assumption ignores basic constraints on male mating success imposed by time, space, and the demands of sperm production, territorial defense, and parental investment. Some models address related constraints by allowing choosy females to remain unmated when acceptable males are not encountered (the “wallflower effect” (De Jong & Sabelis, 1991; Kokko & Mappes, 2005)) or by having offspring per mating decline as a male acquires more matings (Kirkpatrick, 1985; van Doorn et al., 2004). Here, I examine a different consequence of male reproductive limits: successful males become unavailable to subsequent females, directly changing the pool from which females choose. In a natural setting, as female preference spreads and concentrates mating success among preferred males, males are expected to become increasingly unavailable to additional females. Female preference therefore no longer operates in an effectively unlimited mating market, making male availability relevant to the evolutionary fate of preference alleles.

### A gene-centered formulation of the Fisher process

The evolutionary consequences of limits on male mating depend on how declining access to preferred males changes the indirect fitness benefits that allow preference alleles to spread despite their costs. Within the inclusive-fitness framework introduced by Hamilton (1964), the Fisher process can be interpreted as a greenbeard-like process: a preference allele favors copies of itself by directing mating success toward males disproportionately likely to carry the same allele (Faria et al., 2018; Pizzari & Gardner, 2012; Veller et al., 2020). Expressing this logic directly as an allele-frequency recursion makes it possible to trace how access to preferred males affects paternal transmission of the preference allele.

Let *p* denote the frequency of an allele causing female choosiness before selection. For clarity, consider first a haploid model, in which each reproducing individual carries a single copy of an allele at the focal locus. With equal maternal and paternal genetic contributions, the frequency of the preference allele in the next generation is the average of its frequencies among maternal and paternal contributions,

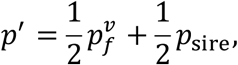

where 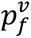 is the frequency of the preference allele among maternal contributions following viability selection against costly female preference, and *p*_sire_ is its frequency among paternal contributions. Because female preference may impose viability or other premating costs, maternal transmission is determined by the allele frequency following selection rather than before selection. The same recursion applies to autosomal diploid inheritance when these quantities are expressed as the marginal frequencies of the focal allele among gene copies transmitted by each sex.

Paternal transmission can be partitioned according to the maternally transmitted allele:

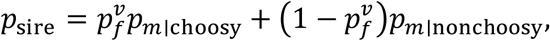

where *p_m_*_∣choosy_ and *p_m_*_∣nonchoosy_ denote the frequencies of the preference allele among paternal allele copies paired with maternally transmitted preference and alternative alleles, respectively (equivalently, the mates of choosy and nonchoosy females in haploid models). These paternal contributions are where male mating limits become relevant: as successful males become unavailable, the changing pool of potential sires can alter the transmission advantage of the preference allele even when female preferences themselves remain unchanged.

To isolate enrichment generated by mate choice from the background allele frequency, the frequency among paternal allele copies paired with preference alleles can be written as the sum of its expectation under random mating and the excess above that expectation:

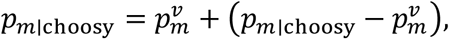

where 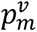 is the frequency of the preference allele among paternal allele copies following viability selection against exaggerated male display. The second term is the excess frequency of the preference allele among paternal copies paired with maternally transmitted preference alleles, relative to a random sample from the paternal allele pool. An analogous expression describes paternal copies paired with maternally transmitted alternative (non-choosy) alleles. Together with the marginal allele frequencies, these absolute enrichment terms completely determine paternal transmission, but they combine two distinct effects: how effectively the mating process pairs preference alleles with paternal copies of the same allele and how much additional enrichment is mathematically possible at the current allele frequency. For example, when the preference allele is already common, even perfect assortment can produce only a small increase above the population average. Absolute enrichment therefore cannot distinguish whether indirect fitness benefits are weak because mating generates little allelic assortment or because little additional enrichment is possible at the current allele frequency. Normalizing enrichment by the distance between the paternal population mean and the focal allele’s genic value yields Grafen’s geometric measure of relatedness (Grafen, 1985):

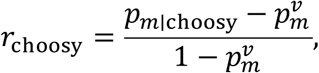

and similarly for the alternative allele,

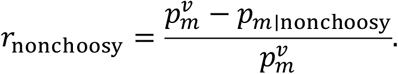

These coefficients describe the allelic relatedness structure generated by mating—each is zero under random pairing and one under maximal positive assortment, while negative values indicate disassortment. Coefficient values depend on the strength of mate choice, genetic associations between preference and display, the availability of preferred males, and, in diploid systems, genotype-to-phenotype relationships such as dominance. Substituting these definitions into the expression for paternal transmission gives

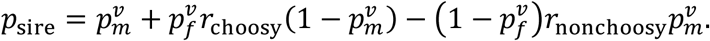

The first term is the baseline contribution expected from the post-selection paternal allele pool. The second term captures the indirect fitness benefit generated when maternally transmitted preference alleles are paired with paternal copies of the same allele more often than expected by chance. The third term captures the corresponding indirect fitness effect for the alternative allele. The indirect component of selection on female preference thus depends not simply on the strength of mate choice, but on the relatedness structure that mate choice generates. Because *r*_choosy_ is calculated from realized maternal-paternal allele pairings rather than preference alone, it depends both on which males females prefer and whether those males are available to sire offspring. All else equal, ecological factors that limit access to preferred males should reduce relatedness and weaken the indirect fitness benefits of mate choice.

The relatedness formulation of the Fisher process immediately suggests why models so often produce unchecked runaway exaggeration. Under the classical assumption that highly successful males can mate without limit, preferred males remain indefinitely available, allowing preference-carrying females to mate consistently with males enriched for the same preference allele and thereby maintain high allelic realized relatedness. When mating capacity is limited, however, preferred males become unavailable as preference spreads and concentrates mating success. This reduction in availability beyond that caused by viability selection should reduce allelic realized relatedness between preference alleles carried by mating partners, throttling the indirect fitness benefits of female preference. The spread of preference alleles is therefore determined by both the strength of female preference and the availability of preferred males.

This gene-centered perspective yields three predictions. First, limits on male mating success should prevent indefinite runaway exaggeration by causing saturated preferred males to become unavailable, reducing realized relatedness until indirect selection no longer favors further escalation. Second, costly female preference and display exaggeration should persist over evolutionary timescales when relatedness remains sufficiently high to offset the costs of preference, but not so high as to drive unchecked runaway evolution. Third, any factor that increases allelic realized relatedness by associating maternally transmitted preference alleles with paternal copies of the same allele carried by preferred males should strengthen sexual selection and promote display exaggeration. Here, I test these predictions using individual-based simulations that track the joint evolution of female preference and male display under different mating limits, preference strengths, recombination rates, and genetic architectures. Together, these analyses show that limits on male mating success can simultaneously sustain costly preference, stabilize exaggerated display, and amplify the additive genetic variation underlying that display.

## Methods

### Individual-based simulations

Individual-based simulations compared limited and unlimited male mating capacity to test how male availability affects the persistence of costly preference, exaggerated display, and genetic variation. Simulations were implemented in R (R Core Team, 2023), with mate choice, mutation, recombination, and offspring production implemented in C++ through Rcpp functions. Simulations followed a population of 50,000 diploid organisms with discrete, non-overlapping generations. Female preference was determined by a single biallelic autosomal locus with alleles denoted choosy and nonchoosy. The locus was inherited by both sexes but expressed only in females and was unlinked to the display loci. Both preference strength and its viability cost were additive: heterozygous females expressed half the preference strength and incurred half the viability cost of choosy homozygotes. Male display was determined by 100 biallelic loci with additive effects. The display of male *i* was

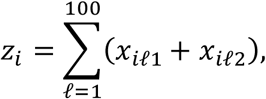

where *x_il_*_1_ and *x_il_*_2_ are the two allelic states at display locus *l*. Display therefore ranged from 0 to 200. There was no environmental component, dominance, or epistasis in display expression. In both autosomal and Z-linked models, display alleles were carried and transmitted by females and males, although their copy number and inheritance differed between the two architectures. Display was expressed and subject to selection only in males.

The choosy allele began at a frequency of 0.05. Initial display-locus allele frequencies were drawn from a beta distribution with shape parameters 0.5 and 0.5, producing a spectrum enriched for rare alleles and approximating the excess of rare variants in empirical allele-frequency spectra. The allele coded 1 was assigned the sampled frequency at half the loci and its complement at the other half, starting mean display near the viability optimum of 100 while creating segregating variation across display loci that was initialized in linkage equilibrium. The same initialization was used for autosomal and Z-linked display. Each generation consisted of viability selection followed by mate choice, mutation, and reproduction.

Male display was subject to Gaussian stabilizing viability selection around an optimum *θ* = 100. A male with display *z* survived with probability

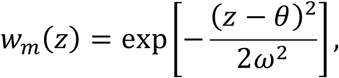

where *ω* = 25 was the width of the Gaussian fitness function and determined the strength of stabilizing selection. This value imposed moderate stabilizing selection: survival was 0.923 for males 10 display units from the optimum and 0.607 for males 25 units from the optimum. Female viability depended on preference genotype, with *s* = 0.01. Females carrying zero, one, or two choosy alleles therefore had relative viabilities 1, 1 − *s*/2, and 1 − *s*, respectively, corresponding to 1, 0.995, and 0.99. Preference had no direct effect on male viability, and display had no direct effect on female viability.

Randomly ordered surviving females next chose mates sequentially. The relative mating weight assigned by female *i* to surviving male *j* was

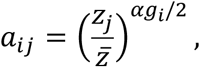

where *z_j_* is the male’s display, 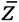 is mean display among surviving males, *α* is preference strength, and *g_i_* is the female’s number of choosy alleles. Males were sampled in proportion to these weights. This is the relative psychophysical preference function of Lande (1981). In treatments with limits on male mating success, a male retained his full display-dependent mating weight until reaching the specified number of matings and was then removed from the pool available to females mating later. In unlimited treatments, males remained available throughout mating.

Display alleles mutated symmetrically between their two states at a rate of 10^−5^ per copy per generation. Preference alleles mutated symmetrically at a rate of 10^−4^. The higher preference-locus mutation rate reduced waiting times for reintroduction following loss of either allele, allowing long-term dynamics to be observed more efficiently. Each mated female produced five offspring, all sired by her assigned mate. From the resulting offspring, 50,000 were sampled at random to form the next generation. A simulation ended and was classified as extinct if no males survived viability selection or if the surviving mated females collectively produced too few offspring to establish the next generation.

### Genetic architectures and parameter sweep

Recombination among display loci was modeled at three levels: free recombination (*ρ* = 0.5), moderate linkage (*ρ* = 0.01), and tight linkage (*ρ* = 0.001). At *ρ* = 0.5, display loci assorted independently, equivalent to placing every display locus on a separate chromosome in the autosomal case. At the two lower rates, the 100 display loci were ordered along a single linkage group, with the specified recombination probability between adjacent loci.

In the autosomal architecture, females and males each carried two homologous sets of display loci, and offspring inherited one set from each parent. Each offspring was assigned male or female with equal probability. The preference locus underwent ordinary Mendelian segregation independently of the display loci. In the Z-linked architecture, females were ZW and males were ZZ. Females therefore carried one set of display alleles and males carried two. Every offspring inherited a paternal Z chromosome. Maternal transmission of the W produced a daughter, whereas transmission of the maternal Z produced a son. Recombination among display loci occurred during paternal gamete production; the maternal Z was transmitted intact because ZW females had no homologous Z chromosome with which it could recombine. The autosomal preference locus segregated independently from both parents as in the autosomal-display model.

Across simulations, preference strength varied from 0 to 50 in increments of 5 and male mating success was limited to 2–10, 100, or unlimited matings per male. The complete parameter grid was evaluated under free recombination, moderate linkage, and tight linkage, with both autosomal and Z-linked display inheritance. Five independent replicate simulations were run for every parameter combination and followed for 50,000 generations unless extinction occurred earlier.

### Summary measures and statistical analysis

Preference allele frequency was recorded in the complete population before viability selection. Mean display, display standard deviation, and additive genetic variance in males were also recorded before viability selection. Because display was entirely additive and had no environmental component, additive genetic variance was calculated as the variance of male display values and includes both variance contributed by individual loci and any covariance arising from associations among loci:

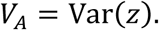

Allelic realized relatedness measures how strongly maternally transmitted alleles are paired through mating with paternal copies of the same allele, relative to the surviving male population. It was calculated each generation after viability selection and mating but before mutation and reproduction. Let 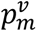 be the frequency of the choosy allele among preference-locus copies carried by surviving males. For each female–sire pair, the numbers of choosy and nonchoosy alleles carried by the female were used to weight the expected frequency of the choosy allele contributed by her mate. This produced the paternal choosy-allele frequencies associated with maternal choosy and nonchoosy allele copies, *p_m_*_∣choosy_ and *p_m_*_∣nonchoosy_, respectively.

Realized relatedness was calculated as

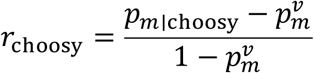

and

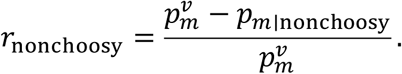

This allele-copy weighting incorporates the choosy and nonchoosy copies carried by heterozygous females and uses the expected allelic contribution of the sire to estimate marginal paternal transmission associated with each maternal allele. The primary analyses used *r*_choosy_, which measures how strongly maternal choosy alleles were paired through mating with paternal copies of the same allele above the surviving-male baseline.

Unless otherwise stated, generation-level quantities were averaged over the final 5,000 generations within each replicate, and the resulting replicate means were averaged across the five simulations to obtain one value for each parameter combination. Reported medians and ranges were calculated from these parameter-combination values. Each α > 0 treatment was matched to an α = 0 treatment having the same male mating limit, recombination rate, and genetic architecture; matched no-preference baselines were calculated as means across the five replicate means. An α > 0 replicate was classified as exhibiting persistent preference if it did not go extinct and its choosy-allele frequency exceeded the corresponding no-preference mutation– selection baseline during at least 95% of the final 5,000 generations. A parameter combination was classified as supporting persistent preference only when this criterion was met in all five replicate simulations.

Standardized display exaggeration was calculated within each replicate by subtracting the matched no-preference (α = 0) mean display from the replicate mean and dividing by the additive genetic standard deviation of the no-preference treatment. Amplification of additive genetic variance was calculated by dividing replicate *V_A_* by the matched no-preference treatment *V_A_*. These replicate-level values were averaged across the five simulations.

Architecture comparisons were conducted using parameter-combination values. For each α > 0 combination of recombination rate, preference strength, and male mating limit, the ratio of Z-linked to autosomal variance amplification was calculated and log-transformed. A linear model with persistence status as its predictor tested whether this architecture ratio differed inside and outside the persistent preference region. Standardized display exaggeration was compared between architectures using a paired *t*-test restricted to parameter combinations that supported persistence under either architecture.

## Results

### Limited mating capacity stabilizes sexual selection

Under unlimited mating capacity and intermediate preference strength, simulated populations exhibited the runaway dynamics predicted by classical Fisherian theory (Fig. 1A). Female preference and male display repeatedly increased to extreme values before collapsing, producing long-term cycles rather than stable exaggeration. During each runaway episode, directional sexual selection drove display-increasing alleles toward fixation as a few highly preferred males monopolized reproductive success, depleting additive genetic variation in display (Fig. 1B). Once this variation was exhausted, mate choice no longer generated the indirect fitness benefits required to maintain costly female preference, causing both preference and display to collapse. Mutation subsequently restored display variation, leading to a new runaway episode.

**Figure 1.**
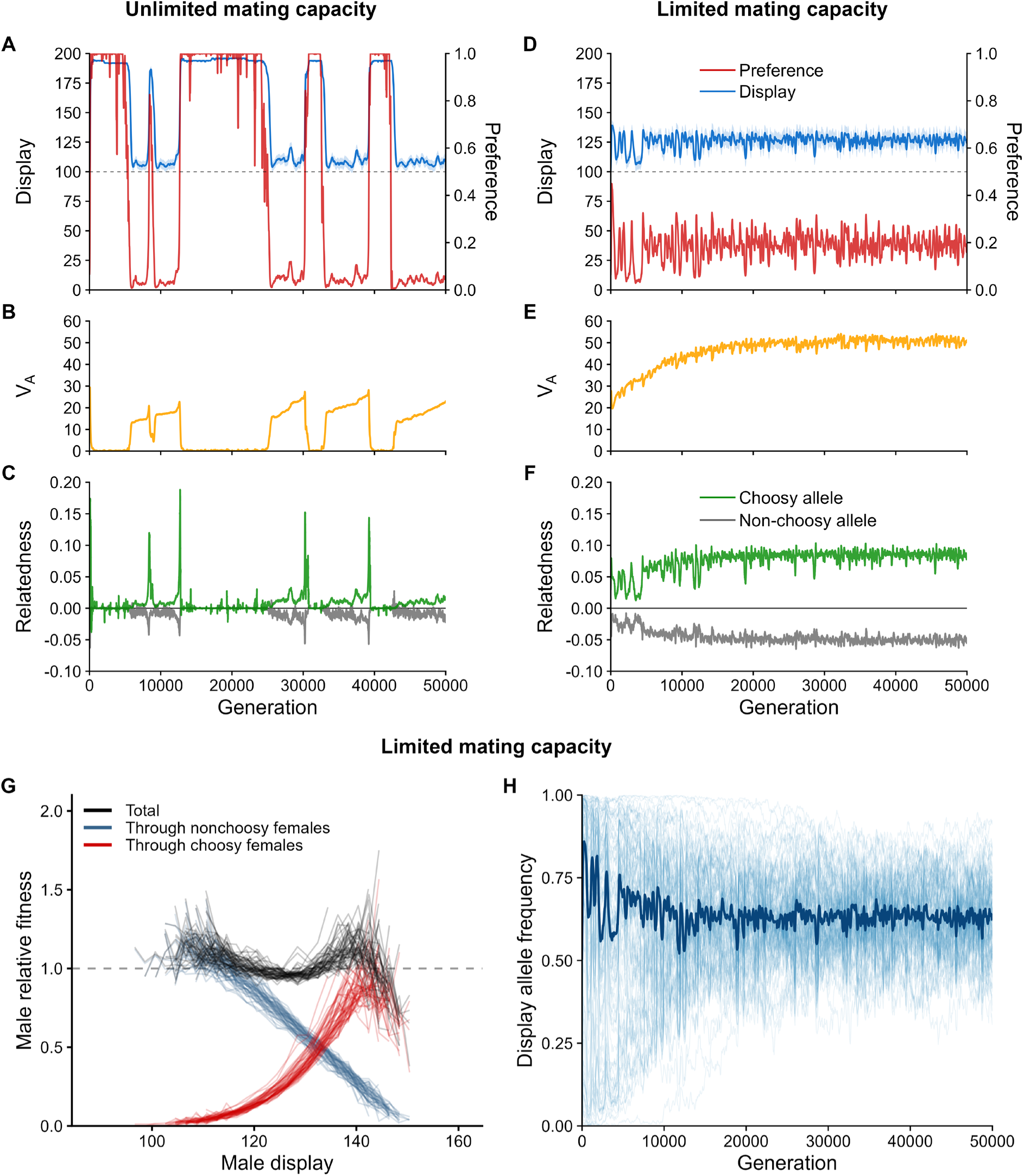
Limits on male mating success stabilize Fisherian dynamics. Evolution of female preference, male display, additive genetic variance in display, and allelic realized relatedness over 50,000 generations under unlimited (A–C) and limited (D–F) male mating capacity. (A, D) Frequency of the female preference allele (red; right axis) and mean male display (blue; left axis, with ±1 SD band). The dashed horizontal line marks the display optimum under stabilizing viability selection. (B, E) Additive genetic variance in male display (*V_A_*). (C, F) Allelic realized relatedness generated by mate choice for the preference allele (*r*_choosy_, green) and alternative allele (*r*_nonchoosy_, gray). (G) Male relative fitness (realized lifetime reproductive success) as a function of display over the last 5,000 generations in the limited-capacity simulation of D-F. Total fitness (black) is partitioned into contributions through nonchoosy (blue) and choosy (red) females. Each curve represents a single generation and curves are shown at 100-generation intervals. (H) Display-increasing allele frequencies at all 100 loci for the limited-capacity simulation of D-F. Thin blue lines show individual loci at 50-generation intervals and the dark blue line shows the median across loci. All results show a single simulation per treatment with preference strength α=35 and free recombination; males could mate a maximum of five times in the limited-capacity simulation shown in D-H.

The mechanism underlying these cycles is evident in the relatedness dynamics (Fig. 1C). During each runaway episode, mate choice increasingly associated maternally transmitted preference alleles with paternal copies of the same allele from favored males, raising allelic realized relatedness and generating the indirect fitness benefits that drive Fisherian sexual selection. As display exaggeration increased, viability selection made the most preferred males progressively rarer. Classical Fisherian models capture this decline in abundance, but because surviving males can mate without limit, rarity does not make them unavailable as mating proceeds. Even a single surviving preferred male remains eligible to mate with every female in the population. Consequently, allelic realized relatedness continues to increase even when the preferred phenotype has become exceedingly uncommon, allowing runaway exaggeration to continue until genetic variation in display is exhausted. Once genetic variation no longer allows choosy females to discriminate among males, allelic realized relatedness collapses, followed by the collapse of costly preference and exaggerated display.

At the same preference strength, limiting male mating capacity fundamentally altered these dynamics (Fig. 1D–F). As preferred males reached their mating limit, their availability to additional choosy females declined, forcing increasing numbers of choosy females to mate with less-preferred males. This prevented the extreme mating skew characteristic of runaway exaggeration and constrained the realized relatedness generated by mating (Fig. 1F), weakening the indirect fitness benefits that drive further evolution of preference and display. Rather than producing repeated cycles of escalation and collapse, this negative feedback led to persistent female preference and stable display exaggeration over tens of thousands of generations (Fig. 1D).

Importantly, stabilization did not require the elimination of sexual selection. Sexual selection remained strong at this equilibrium: in the representative stabilized run, mean display over the final 5,000 generations was 11.2 additive-genetic SDs above its matched no-preference baseline, and a male expressing that mean retained only 57% of the viability of a male at the viability optimum (Fig. 1D). Preference and display continued to fluctuate as changing male availability altered realized relatedness and the indirect fitness benefits of choice, maintaining costly preference and display exaggeration over evolutionary time rather than leading to escalation without bound or permanent collapse.

Within this regime, the combined effects of viability and mating success generated disruptive selection on male display (Fig. 1G). Males with relatively low displays benefited from greater survival and opportunities to reproduce with nonchoosy females, whereas more exaggerated males compensated for lower survival through greater mating success with choosy females. Intermediate displays yielded lower lifetime reproductive success than displays on either side of the fitness minimum. This disruptive selection favored the maintenance of display variation even as the relative advantages of lower and higher displays fluctuated. Display-locus allele frequencies converged toward intermediate values and remained broadly polymorphic (Fig. 1H), sustaining the elevated additive genetic variance observed under limited mating capacity.

### Relatedness links mating constraints to persistent sexual selection

A parameter sweep across male mating capacity, female preference strength, and recombination rate revealed a broad region of parameter space supporting persistent female preference and stable display exaggeration (Fig. 2). Black outlines identify parameter combinations in which female preference remained above the mutation-selection baseline expected in the absence of female choice (α=0) for at least 95% of the final 5,000 generations in every replicate simulation. These combinations supported persistent female preference and were also characterized by stable display exaggeration. When limits on male mating success were sufficiently strong, costly female preference and exaggerated male display were sustained across a range of preference strengths despite ongoing viability selection against both choosy females and high-display males. Across all 49 parameter combinations supporting persistent preference, mean display exceeded the matched no-preference (α=0) baseline maintained by stabilizing selection on display alone by a median of 12.5 SDs (range 4.4–16.5).

**Figure 2.**
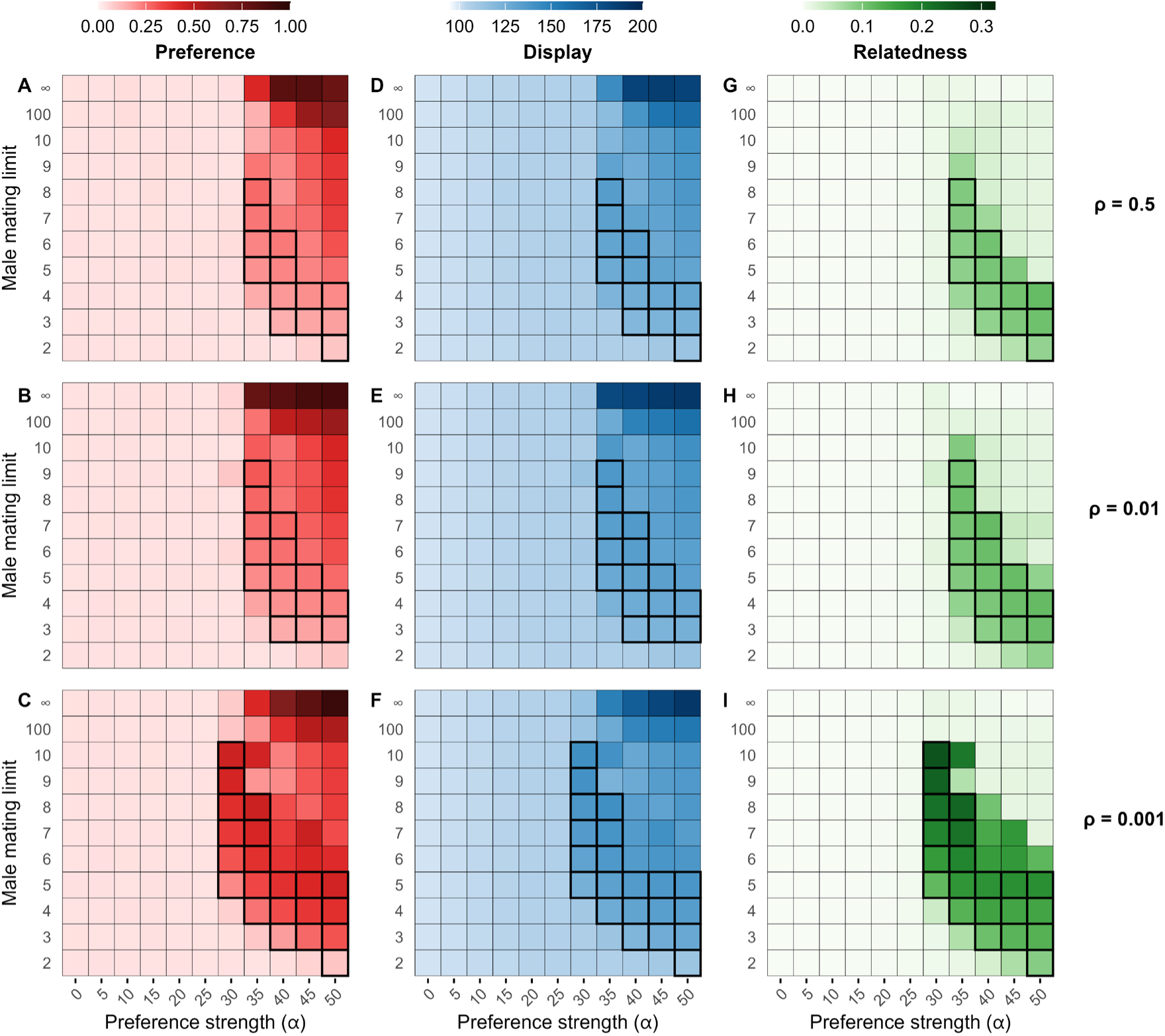
Genetic linkage broadens the conditions for stable preference and display. Effects of preference strength (*α*), the maximum number of matings per male, and recombination among display loci on female preference (A–C), male display (D–F), and allelic realized relatedness generated by mate choice (*r*_choosy_; G–I). Rows show results under free recombination (*ρ* = 0.5), moderate linkage (*ρ* = 0.01), and tight linkage (*ρ* = 0.001). Values were first averaged over the final 5,000 generations of each simulation and then across simulations (n = 5 per parameter combination). Black outlines identify parameter combinations in which female preference remained above the matched α=0 baseline—with the same male mating limit and recombination rate—for at least 95% of the final 5,000 generations in all five replicate simulations. As recombination declined, progressively weaker preference was sufficient for persistence of female preference, expanding the region of parameter space supporting stable sexual selection.

At low preference strengths, female preference either failed to spread or spread only transiently (Fig. 2A–C). The indirect fitness benefits generated by mate choice were insufficient to offset the cost of choosiness, even though viability selection against male display remained weak because display values were close to the naturally selected optimum. Increasing preference strength promoted both female preference and display exaggeration, but these traits persisted only within a restricted region of parameter space (outlined in Fig. 2). Outside this region, particularly when male mating limits were weak or absent, high mean preference and display often reflected final sampling windows that captured the exaggerated phase of an ongoing runaway cycle. Extreme display accompanied by low allelic realized relatedness marked late-stage runaway peaks, after display variation had been exhausted and mate choice could no longer direct substantial indirect fitness benefits toward additional copies of the preference allele. These populations were approaching another collapse of preference and display.

Reducing recombination expanded the region of parameter space supporting persistent preference and display exaggeration (Fig. 2). As recombination declined, the number of persistent parameter combinations increased from 13 under free recombination to 15 under moderate linkage and 21 under tight linkage, with progressively weaker preference sufficient for persistence. Reduced recombination preserved multilocus associations among display alleles that would otherwise be continually broken apart, maintaining high-display genetic backgrounds that mate choice could associate with preference alleles and thereby sustain elevated realized relatedness. Across all three regimes, persistence closely coincided with elevated relatedness (Fig. 2G–I), consistent with the predicted role of relatedness in maintaining costly preference and exaggerated display.

### Sexual selection maintains elevated genetic variation

Stable sexual selection consistently amplified the genetic variation underlying display. Across a broad range of parameter combinations, additive genetic variance in male display substantially exceeded that expected under mutation-selection balance in the absence of female preference (Fig. 3). For parameter combinations supporting persistent preference and display, additive genetic variance was universally elevated, ranging from 7.9- to 32.2-fold higher than in matched no-preference treatments. Median increases were 9.0-fold under free recombination (Fig. 3A), 9.4-fold under moderate linkage (Fig. 3B), and 21.3-fold under tight linkage (Fig. 3C). Genetic variance was generally greatest where sexual selection remained strong but preferred males could not monopolize reproduction. Reduced recombination further amplified this effect by preserving display haplotypes, producing exceptionally high additive genetic variance in some regions of parameter space. By maintaining the display variation that allows female choice to continue generating indirect fitness benefits, these dynamics reverse the classical expectation at the heart of the lek paradox: persistent female choice amplifies, rather than depletes, the genetic variation on which choice depends.

**Figure 3.**
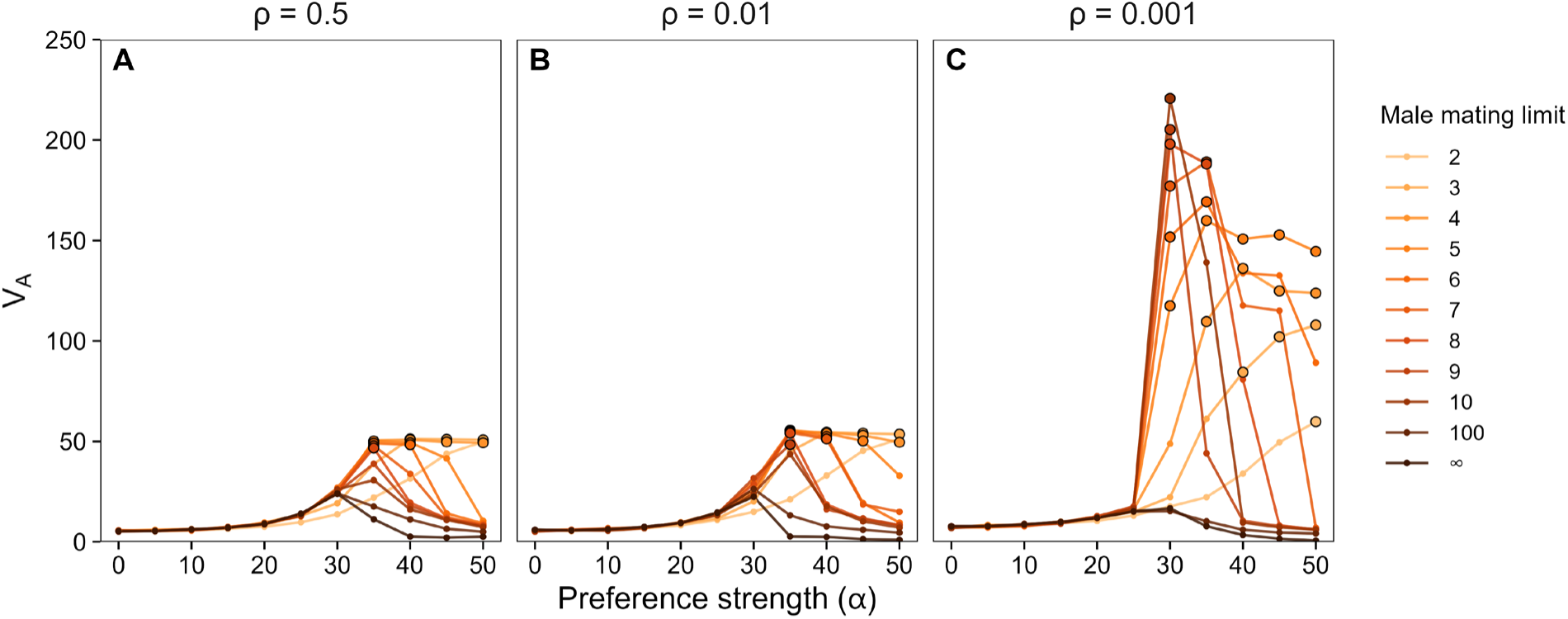
Sexual selection amplifies genetic variation. (A–C) Additive genetic variance in male display (*V_A_*) across preference strengths (*α*) and male mating limits under free recombination (*ρ* = 0.5, A), moderate linkage (*ρ* = 0.01, B), and tight linkage (*ρ* = 0.001, C). Colors indicate the maximum number of matings per male. Points show values averaged over the final 5,000 generations within each replicate and then across the five replicate simulations. Black outlines identify parameter combinations for which all five replicates met the persistence criterion. Female preference amplified additive genetic variance above matched no-preference (α = 0) baselines most strongly under mating limits that maintained elevated allelic relatedness, with the magnitude of amplification generally increasing as linkage among display loci increased.

### Z linkage of display loci expands the parameter space supporting stable exaggeration

Sex linkage can alter the genetic association between female preference and male display that drives Fisherian sexual selection (Kirkpatrick & Hall, 2004). Because female-heterogametic (ZW) sex determination characterizes birds and many other taxa with elaborate male ornamentation, the Z chromosome is a particularly relevant location for display loci. If Z linkage increases the realized allelic relatedness generated by mate choice, it should expand the parameter space supporting persistent female preference and display exaggeration. To test this prediction, I repeated the parameter-space analysis with all 100 display loci located on the Z chromosome rather than on autosomes. To isolate the effect of chromosome identity, the preference locus remained autosomal in both architectures.

Z linkage expanded the region of parameter space supporting persistent female preference and display exaggeration from 15 to 27 combinations (Fig. 4, shown only for ρ=0.01; Z-linked results across all three recombination regimes are shown in Fig. S1). Parameter combinations that failed to sustain exaggerated display when display loci were autosomal frequently supported persistent exaggeration when those loci were moved to the Z chromosome (Fig. 4A–D). Across the 27 parameter combinations supporting persistence under at least one genetic architecture, mean display was elevated above the matched no-preference baseline by a median of 11.6 SDs (range 2.6–16.5) under autosomal inheritance and 16.1 SDs (range 6.9–20.8) under Z-linked inheritance. Z linkage produced greater standardized exaggeration in all 27 combinations, with a median increase of 29% relative to autosomal inheritance. Across these matched combinations, standardized display exaggeration was on average 5.3 SD units greater under Z-linked than autosomal inheritance (95% CI 3.7–7.0, *t*_26_ = 6.68, *P* < 0.001). Z linkage also produced a parallel increase in realized relatedness (Fig. 4E,F).

**Figure 4.**
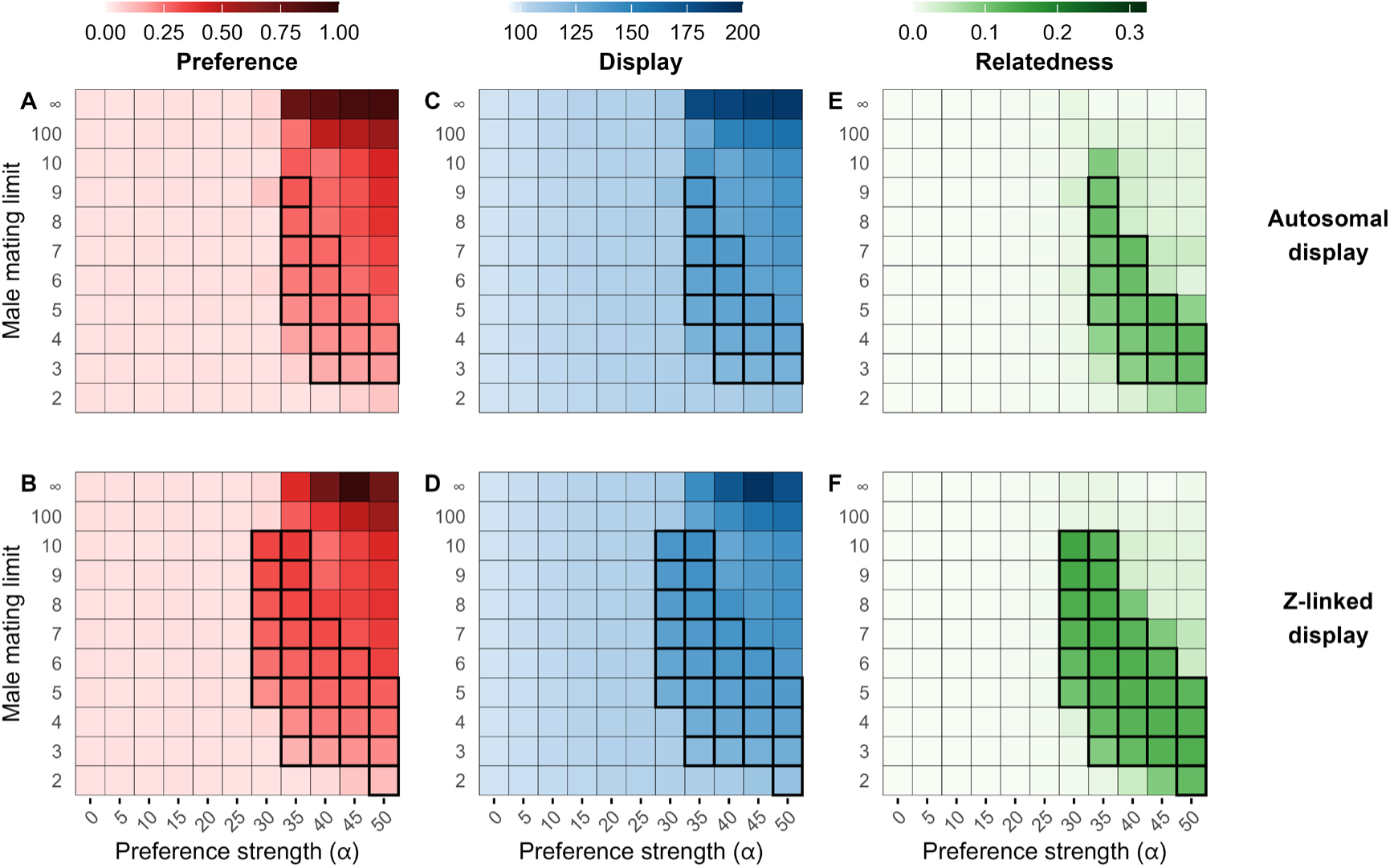
Z-linked display broadens the conditions for stable preference and display. Effects of preference strength (α), the maximum number of matings per male, and the genetic architecture of display on female preference (A–B), male display (C–D), and allelic realized relatedness generated by mate choice (E–F). Rows show results when display loci were autosomal or Z-linked. Recombination among display loci was moderate (ρ=0.01) in both treatments. Black outlines identify parameter combinations in which female preference remained above the matched α = 0 baseline—with the same male mating limit and display architecture—for at least 95% of the final 5,000 generations in all five replicate simulations. Relative to autosomal display, Z linkage maintained elevated allelic realized relatedness under weaker preference and less restrictive mating limits, expanding the region of parameter space supporting persistent female preference and stable display exaggeration.

Across all three recombination regimes, Z linkage likewise strengthened the amplification of additive genetic variance in display (Fig. S2). Amplification was greater under Z-linked than autosomal inheritance in 73/74 (99%) of parameter combinations supporting persistence under at least one architecture. Within this region, the geometric-mean ratio of Z-linked to autosomal variance amplification was 1.63 (95% CI 1.52–1.74), compared with 1.10 outside the persistence region (95% CI 1.06–1.14). The ratio of Z-linked to autosomal amplification of genetic variance was therefore 1.48 times as large where preference persisted than where it did not (95% CI 1.37– 1.60, *t*_328_ = 9.94, *P* < 0.001).

These effects arise from the asymmetric transmission of Z-linked display alleles under female heterogamety. When a choosy female mates with a highly ornamented male, daughters that inherit her preference allele carry it together with a Z chromosome inherited from the preferred father. Because ZW daughters have only this single Z, the paternal display haplotype is not paired with a maternal Z, does not recombine, and is transmitted intact to every son. Under autosomal inheritance, paternal display alleles are paired with maternal homologs that need not have high breeding values, can be separated by recombination, and are each transmitted to only half of a daughter’s sons on average. Z linkage therefore transmits display alleles from preferred fathers more reliably through daughters carrying maternally inherited preference alleles, strengthening preference–display associations across generations. Highly ornamented males consequently become more likely to carry the preference allele, allowing choosy females to direct more paternal transmission toward additional copies of their own allele. Ecological limits on male mating success and Z linkage therefore act through complementary routes: the former limits access to preferred males, stabilizing costly preference and display, whereas the latter strengthens preference–display associations, allowing persistent exaggeration across more conditions and at higher levels. By jointly regulating realized relatedness, these mechanisms may help explain the abundance of exaggerated male displays in female-heterogametic taxa.

## Discussion

The central result of this study is that limits on male mating success generate negative feedback that allows costly female preference and exaggerated male display to persist across a broad region of parameter space without runaway escalation. When successful males can mate without limit, preference and display repeatedly escalate, exhaust the display variation sustaining the Fisher process, and collapse. Restoring the biologically unavoidable constraint that is abstracted away by unlimited-mating assumptions alters these dynamics. By reducing allelic realized relatedness as preferred males become unavailable, these limits interrupt runaway evolution while preserving costly preference, exaggerated display, and the genetic variation on which female choice depends. Natural display exaggeration can be truly spectacular: male tail ornaments in three avian families average roughly 30 phenotypic SDs beyond the corresponding unexaggerated traits (Alatalo et al., 1988). This scale has served as a benchmark for evaluating the capacity of sexual-selection models to generate extreme exaggeration (Houle & Kondrashov, 2002; Kirkpatrick, 1996). Even with a deliberately simplified 100-locus display architecture, no contribution of environmental variation, and a 1% viability cost of female preference, stable Fisherian sexual selection produced shifts above matched no-preference baselines of up to 16.5 SDs with autosomal display (20.8 SDs with Z-linked display), approaching this extraordinary empirical scale.

These results also suggest a different way of understanding the Fisher process itself. Fisherian sexual selection is typically discussed in terms of linkage disequilibrium, genetic correlations, or runaway dynamics. Although allelic realized relatedness can be expressed as a genetic association, it is not simply another name for the conventional genetic covariance between preference and display. Preference–display genetic covariance describes the association between breeding values for the two traits within the population. Allelic realized relatedness instead measures the enrichment of paternal preference alleles paired through mating with maternally transmitted preference alleles, relative to the surviving male pool. This enrichment depends on both existing genetic associations and the allocation of matings. Consequently, populations with identical genetic structure before mating can generate different realized relatedness when limits on male mating capacity alter access to preferred males.

The mathematical framework developed here places this process naturally within Hamilton’s theory of inclusive fitness (Hamilton, 1964), extending the greenbeard-like interpretation of Fisherian sexual selection (Faria et al., 2018; Pizzari & Gardner, 2012; Veller et al., 2020). Viewed through this inclusive-fitness lens, sexual reproduction is necessarily a duet: it requires a partner, and mate choice determines what that partnership provides. In the sexual selection process considered here, that return arises neither from direct benefits nor from genome-wide genetic quality. Instead, preference-carrying females incur a fitness cost while directing reproductive success toward males more likely than chance to carry the same preference allele. The fate of the preference allele depends on the same balance between costs and relatedness-weighted indirect benefits that underlies inclusive-fitness theory more generally. So long as allelic realized relatedness remains positive, the display favored by choosy females is an honest indicator of the indirect genetic benefits available to choosy females. This form of honest signaling requires no association between display and condition or genome-wide genetic quality but instead arises from the preference–display association generated by mate choice. Evolution of female preference under the Fisher process is sometimes described as non-adaptive, but this conflates organismal costs with the fate of the alleles that generate them. Sexual selection can impose substantial costs on choosy females yet remain adaptive—preference alleles recoup direct costs paid by female carriers by increasing the reproductive success of copies carried by males.

Direct-benefit and good-genes mechanisms depend on associations between mate choice and external benefits or genetic quality, whereas Fisherian sexual selection generates its preference– display associations through mate choice itself (Kokko et al., 2002). Limits on male availability then regulate whether the resulting indirect benefits can maintain costly preference without runaway evolution. This feedback from male availability may help reconcile disagreement over the importance of Fisherian sexual selection. Previous models have maintained costly preference and display exaggeration by introducing processes that counter their loss or runaway escalation. One invokes directionally biased mutation, which continually introduces alleles opposing exaggeration (Pomiankowski et al., 1991). Another assumes that the cost of choosing a preferred male declines as that male type becomes common (Kokko et al., 2015). Limits on male mating success operate differently. Stabilization arises from a self-limiting consequence of mate choice rather than from a separate force imposed to counter its evolutionary dynamics. In this sense, the Fisher process may provide the simplest general explanation for the evolution of sexual displays.

The amplification of display variation under limited male mating capacity provides a resolution to the lek paradox—the puzzle of how female choice can sustain indirect benefits when sexual selection is expected to erode the genetic variation on which they depend (Borgia, 1979; Kirkpatrick & Ryan, 1991; Rowe & Houle, 1996)—through disruptive selection generated by the interaction of mate choice and viability selection. By maintaining choosy and nonchoosy females together, mating limits preserve alternative routes to male reproductive success: highly exaggerated males gain mating advantages from choosy females, whereas less-exaggerated males benefit from greater viability and opportunities to mate with nonchoosy females. The resulting selection maintains polymorphism at display loci, sustaining the genetic variation on which the indirect benefits of choice depend. In the simulations, this mechanism amplified additive genetic variance as much as 32-fold with autosomal display (44-fold with Z-linked display) relative to matched no-preference treatments. Recent theory shows that mate choice can increase genetic variation through differences among female preferences and genetic associations between mating partners (Xu & Servedio, 2026). Importantly, the present results show how feedback through declining male availability stabilizes Fisherian dynamics, allowing elevated variation to persist alongside costly preference and exaggerated display without requiring mutational biases (Pomiankowski et al., 1991) or the evolution of condition-dependent signaling (Rowe & Houle, 1996). This mechanism also leads to a counterintuitive prediction: if mate choice were to disappear, as in experimental manipulations removing sexual selection, mean nonsexual fitness should increase as costly preference and exaggeration recede, while standing genetic variation for display simultaneously declines. Female preference was controlled by a single biallelic locus in these simulations. Whether mating limits also maintain elevated additive genetic variance in display when preference has a polygenic quantitative basis remains to be tested.

The coexistence of choosy and nonchoosy females at equilibrium may also facilitate the evolution of reproductive isolation through female choice. Genetic associations between preference and display create differences in the multilocus display backgrounds carried by these alternative female types. Given the disruptive male fitness landscape (Fig. 1G), crosses between alternative preference–display backgrounds produce sons with intermediate displays that incur viability costs yet lack the mating advantages of highly exaggerated males—the worst of both worlds. Selection against such sons would favor mating mechanisms that reduce cross-type mating, preserving alternative genetic associations and strengthening assortative mating and reproductive isolation (Kirkpatrick & Ravigné, 2002; Servedio & Noor, 2003). The simulations also showed that Z-linked display variation sustained costly preference across a broader range of conditions and permitted exaggeration under weaker preference. These effects arise from more faithful transmission of preference–display associations across generations and consequently greater allelic realized relatedness between mating partners, suggesting that genetic architectures that preserve these associations may facilitate speciation.

The ant and the peacock embody two of the most profound evolutionary problems confronted by Darwin—the evolution of costly social behavior and extravagant sexual traits (Cronin, 1991). Although social evolution and sexual selection have generally been treated as distinct evolutionary phenomena, both can be understood through the same gene-level logic: an allele can spread when a cost expressed through one carrier increases the reproductive success of copies of that allele in others. In Fisherian sexual selection, this logic is realized when a costly female preference directs mating success toward males carrying the same preference allele. Natural limits on male mating success provide a stabilizing feedback: as preference spreads, preferred males become increasingly unavailable, weakening the indirect benefits of further escalation. With this feedback, the same Fisher process that can drive runaway evolution instead readily sustains costly preference and exaggerated display over evolutionary timescales. Stable sexual selection may therefore be expected rather than exceptional, helping explain both the persistence of elaborate sexual displays and the extraordinary diversity of sexual ornaments observed throughout nature.

## Data and code availability

Simulation code and data available upon publication.

## Author contributions

BH conceived of the project, designed, developed, and executed the simulations, analyzed the results, and wrote the paper.

## Acknowledgements

I thank T.J. Kawecki and F. Hollis for helpful discussion and suggestions on earlier versions of the manuscript. Simulations were performed on the USC Hyperion cluster.

---

**Fig. S1.**
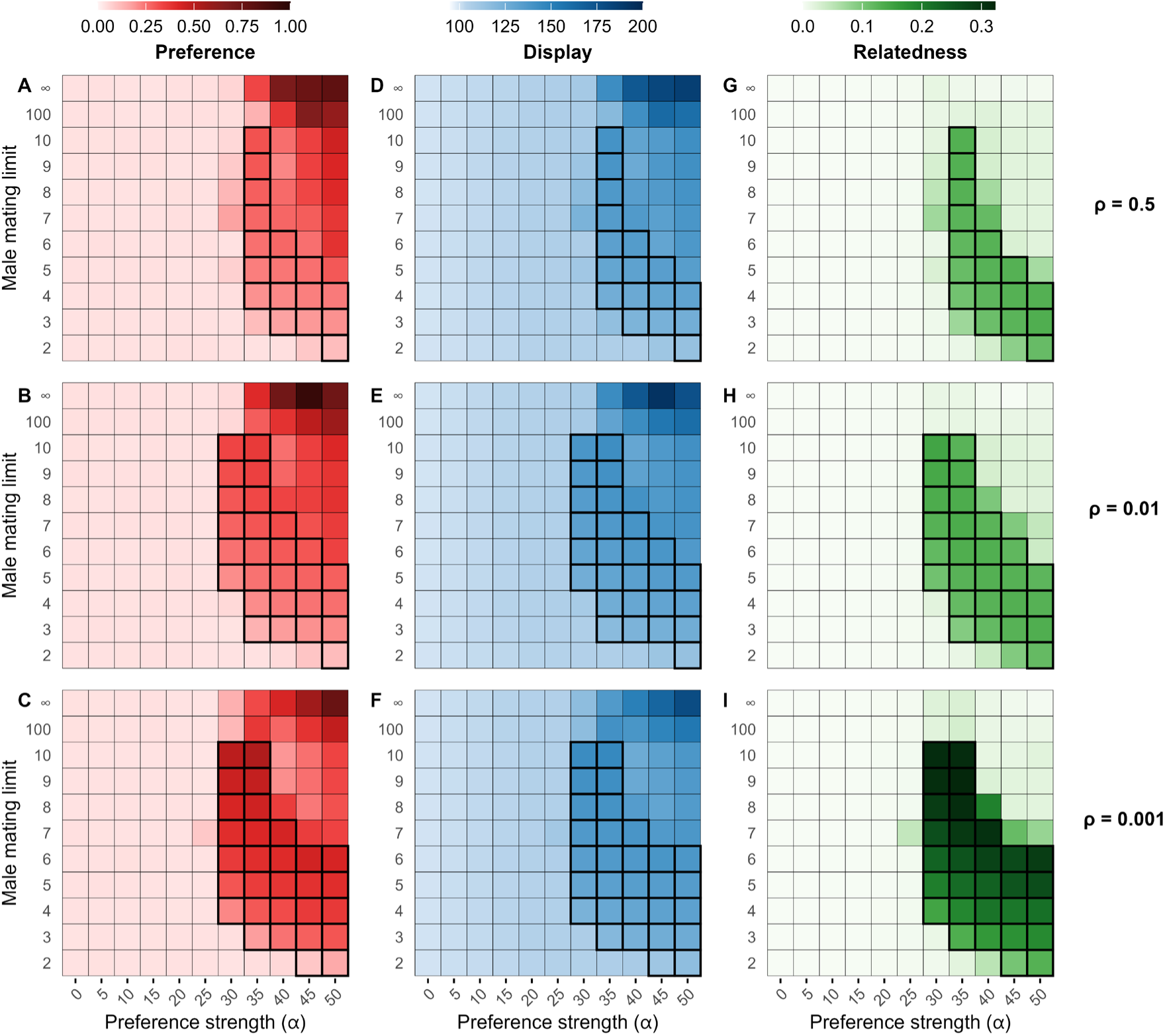
Effects of preference strength (*α*), the maximum number of matings per male, and recombination among display loci on female preference (A–C), male display (D–F), and allelic realized relatedness generated by mate choice (*r*_choosy_; G–I) when display loci are Z-linked. Rows show results under free recombination (*ρ* = 0.5), moderate linkage (*ρ* = 0.01), and tight linkage (*ρ* = 0.001). Values were first averaged over the final 5,000 generations of each simulation and then across simulations (n = 5 per parameter combination). Black outlines identify parameter combinations in which female preference remained above the matched α = 0 baseline—with the same male mating limit and recombination rate—for at least 95% of the final 5,000 generations in all five replicate simulations.

**Fig. S2.**
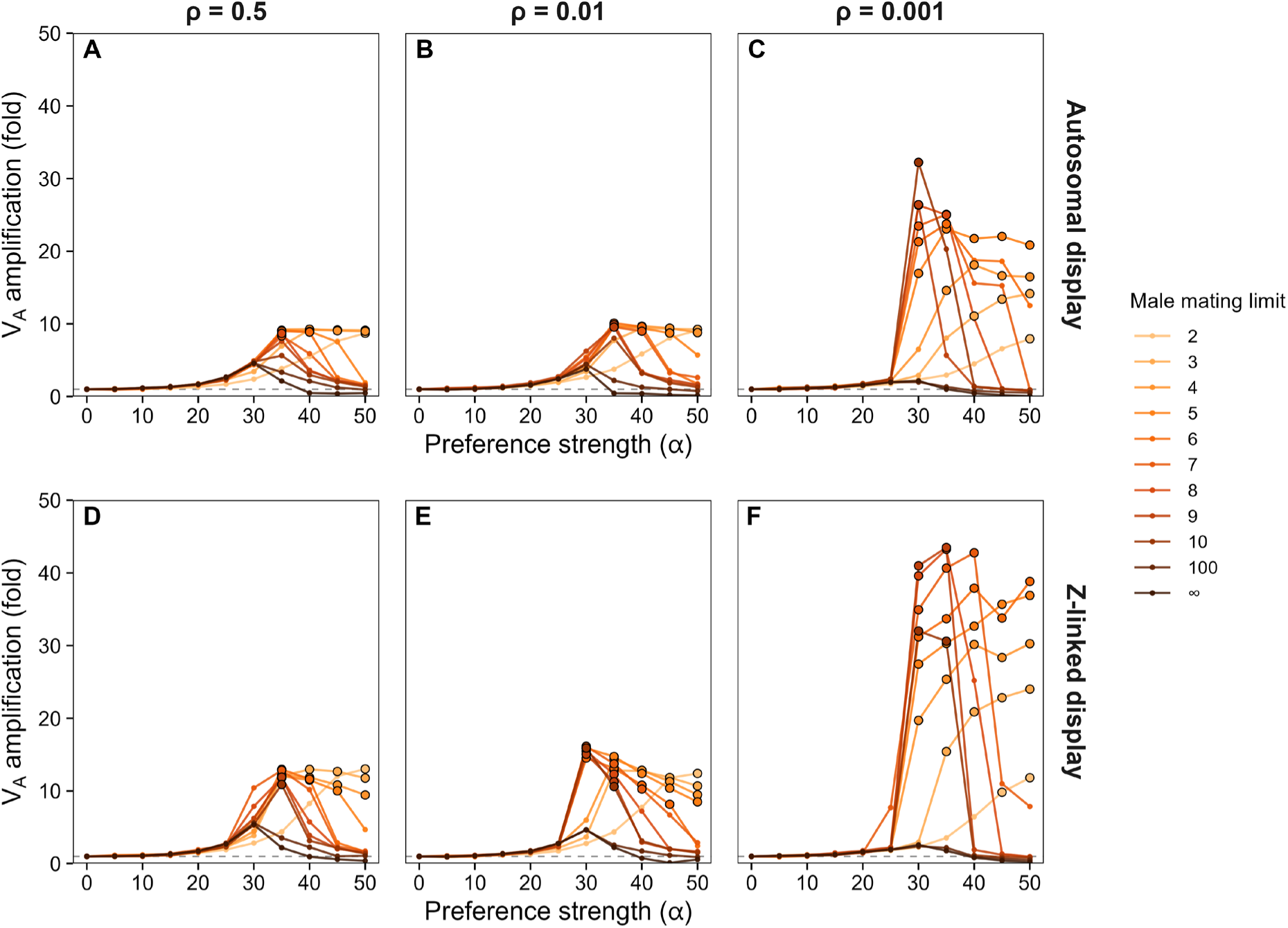
Z linkage of male display enhances the amplification of genetic variation. Fold amplification of additive genetic variance in male display (*V_A_*) across preference strengths (*α*) and male mating limits under free recombination (*ρ* = 0.5), moderate linkage (*ρ* = 0.01), and tight linkage (*ρ* = 0.001), shown for autosomal (A-C) and Z-linked (D-F) display inheritance. Colors indicate the maximum number of matings per male. Within each replicate, *V_A_* was averaged over the final 5,000 generations and divided by the matched no-preference baseline for the same display architecture, male mating limit, and recombination rate. Points show mean amplification across the five replicate simulations. The dashed horizontal line indicates no amplification and black outlines identify parameter combinations where all five replicates met the persistence criterion.

